# Assessing the Functional-Structural Reorganization of Connector Hubs in Temporal Lobe Epilepsy

**DOI:** 10.64898/2026.09.25.754445

**Authors:** Arielle Dascal, Édouard Delaire, Alexander Ngo, Judy Chen, Ella Sahlas, Thaera Arafat, Chifaou Abdallah, Mahdi Mobarak-Abadi, Jessica Royer, Raul Rodriguez Cruces, Neda Bernasconi, Andrea Bernasconi, Raluca Pana, Kangjoo Lee, Boris C. Bernhardt, Christophe Grova

## Abstract

Recent years have seen a rise in investigations on the organizational principles of brain networks in healthy and pathological conditions. Temporal lobe epilepsy (TLE) is a compelling case as its focal pathology is accompanied by widespread network disruptions. However, how TLE affects functional connector hubs, regions supporting long-range and inter-network communication, remain poorly understood. We mapped connector hub reorganization in patients with TLE relative to healthy controls. Patients exhibited widespread connector hub reductions in the ipsilateral occipital, cingulate, precuneus and hippocampal regions as well as the bilateral pallidum. Hub reductions occurred alongside the emergence of new hubs in contralateral regions. We furthermore observed strong associations between hub reductions and microstructural alterations in subcortical structures and the hippocampus. These findings demonstrate widespread, asymmetric connector hub reorganization extending beyond the epileptogenic temporal lobe, revealing distinct patterns of network disruption and brain-wide reorganization.

## 1 Introduction

Epilepsy is a neurological disorder that affects nearly 50 million people worldwide (World Health Organization, 2024). Among patients with drug-resistant epilepsy, temporal lobe epilepsy (TLE) accounts for nearly two-thirds of refractory cases (Asadi-Pooya et al., 2017; Chowdhury et al., 2021). Temporal lobe epilepsy (TLE), a common pharmaco-resistant epilepsy in adults, is characterized by seizures originating from the hippocampus (Engel Jr., 2001). Although surgical intervention can provide seizure freedom for many patients, approximately half experience seizure recurrence following surgery (Tisi et al., 2011; Krucoff et al., 2017). These outcomes suggest that the underlying mechanisms involved in seizure generation and propagation remain misunderstood, motivating the search for imaging biomarkers that may improve surgical planning and outcome prediction.

Increasing evidence suggests that epilepsy should be conceptualized as a network disorder, characterized by widespread alterations extending beyond the hypothesized seizure onset zone and involving distributed cortical and subcortical systems (Spencer, 2002; Bartolomei et al., 2017; Royer et al., 2022). Multimodal neuroimaging studies have demonstrated extensive functional and structural reorganization in TLE both within and beyond mesio-temporal regions. Abnormalities in functional connectivity have been observed using resting state magnetic resonance imaging (rs-fMRI), (Liao et al., 2010; Pittau et al., 2012; Bernhardt et al., 2016; Lee et al., 2018), and electrophysiology (Englot et al., 2015; Aydin et al., 2020; Pourmotabbed et al., 2020; Roehri et al., 2026), whereas structural alternations have been estimated through grey matter morphology (Bernhardt et al., 2009; Whelan et al., 2018), vasculature (Ngo et al., 2024) and white matter microstructure (Concha et al., 2009; Bernhardt et al., 2016; Hatton et al., 2020; Gleichgerrcht et al., 2021). These findings have motivated increasing interest in characterizing large-scale network brain reorganization in epilepsy.

Resting-state functional magnetic resonance imaging (rs-fMRI) enables investigation of large-scale brain organization through spontaneous fluctuations in the blood-oxygen-level-dependent (BOLD) signal, allowing characterization of resting state networks (RSNs) (Fox and Raichle, 2007; Smith et al., 2009; Biswal et al., 2010). This grants insight into large-scale brain organization, which is characterized by spatially unique and temporally correlated networks spanning cortical and subcortical systems (Yeo et al., 2011; Seitzman et al., 2019). In the case of temporal lobe epilepsy, changes in functional connectivity derived from rs-fMRI have commonly been studied using both seed-based correlation and graph theoretical approaches. Typically selecting regions associated with the hypothesized seizure onset zone as a region of interest, seed-based studies have demonstrated increased within-network connectivity in sensory networks (Zhang et al., 2009), decreased connectivity between the anterior and posterior structures of the default-mode network (Haneef et al., 2012; Zhou et al., 2021) and decreased connectivity between the ipsilateral mesial structures with contralateral mesial and bilateral default mode structures (Pittau et al., 2012; McCormick et al., 2013; Bernhardt et al., 2016). In contrast, graph theory enables whole brain nodal analysis, where reduced degree in the ipsilateral epileptogenic structures and default mode regions have been linked to reduced overall network efficiency (McCormick et al., 2013; Ofer et al., 2019; Zhou et al., 2025). Particular interest has been directed toward connector hubs, key regions exhibiting several connections between RSNs (Bertolero et al., 2018), which play a key role in long-range and inter-network communication (Van Den Heuvel and Sporns, 2013). Previous work has demonstrated altered hub organization in both generalized (Zhang et al., 2011) and focal (Lee et al., 2018; Qu et al., 2026) epilepsies. Epilepsy-related changes in hub connectivity have been found to be associated with prolonged seizure duration (Zhou et al., 2025), increased seizure frequency (Douw et al., 2015), cognitive impairment (Roger et al., 2020), and have been shown to help improve the prediction of surgical outcomes (He et al., 2017; DeSalvo et al., 2020; Akbarian et al., 2024; Karpychev et al., 2026). For a comprehensive review see Royer et al., 2022a.

While these methodologies are widely used in brain connectivity analyses, they may not fully capture the complexity of whole-brain functional organization. Seed-based and graph theoretical approaches often assume non-overlapping network architecture, are sensitive to multicollinearity arising from correlated signals, and provide limited information regarding the specific network composition of highly inter-connected hub regions (Lee et al., 2016). However, growing evidence suggests that brain regions can participate in multiple functional systems simultaneously, supporting a flexible and overlapping organization of underlying RSNs (Najafi et al., 2016; Xu et al., 2016). To address these limitations, we developed SParsity-based Analysis of Reliable *k-*hubness (SPARK), a data-driven sparse modeling framework that highlights overlapping structure of functional networks and estimates connector hubs at the voxel level (Lee et al., 2016). SPARK offers the unique ability to identify the many RSNs associated with each hub, while quantifying hubness changes occurring in pathological conditions, when compared to normative healthy data, using the hub disruption and emergence indices (Achard et al., 2012; Lee et al., 2018).

Moreover, extensive research using diffusion weighted imaging (DWI) allowed the identification of structural alterations associated with epilepsy. Mature and intact white matter tracts allow for fast and efficient transmission of information in the brain, whereas epilepsy-related changes are associated with cognitive dysfunction (McDonald et al., 2014; Rodríguez-Cruces and Concha, 2015; Rodríguez-Cruces et al., 2020). Using fractional anisotropy (FA) and apparent diffusion coefficient (ADC) maps derived from DWI data (Alexander et al., 2007), we can observe microstructural changes in the white matter before they can be identified using typical T1 weighted imaging (Zhang et al., 2025). While changes in TLE relative to controls involving reductions in FA and increases in ADC are typically widespread and bilateral, the strongest effects are observed in the ipsilateral temporo-limbic systems that are proximal to the seizure generating region (Otte et al., 2012; Campos et al., 2015; Bernhardt et al., 2016; Lin et al., 2020).

Understanding the complex relationship between structure and function in the human brain can provide insights into information flow in the brain, and especially connector hubs. Structural connectivity provides information about the anatomical framework that constrains and shapes patterns of functional connectivity across the brain (Messé et al., 2014; Ali et al., 2025). In epilepsy, decreases in functional connectivity have typically been associated with the expected changes of increased ADC and decreased FA in regions associated with the epileptogenic focus (Liu et al., 2016). Previous work has indicated that the strongest decreases in functional connectivity in patients with temporal lobe epilepsy were associated with the connector hub nodes, which exhibited a moderate overlap with microstructural changes of FA and ADC (Larivière et al., 2020; Xie et al., 2025). Previously, our group also performed hub mapping using SPARK on a limited cohort of mTLE patients to investigate alterations associated with the bilateral hippocampi, observing significant hub disruption in ipsilateral mesial temporal regions and within the default mode network (Lee et al., 2018). In the present study, we extend upon these prior works by investigating vertex-level patterns of connector hub reorganization across the cortex and subcortex in TLE. We also assess corresponding alterations in the anatomical substrate associated with identified hubs using DWI. Furthermore, leveraging the uniqueness of the SPARK method to extract specific RSNs with their spatial overlap for each subject, we evaluate the distribution of networks contributing to the observed hub disruption associated with epileptic pathology. Moreover, to quantify the rate at which connector hubs are losing connectivity, we propose a modified version of the hub disruption index (Achard et al., 2012; Lee et al., 2018), specifically assessing hub disruption in patients when compared to healthy controls. Finally, considering a subgroup of patients who underwent surgical resection and who thereafter became seizure free (Engel IA status) after one year of follow up (Epilepsy (ILAE) 1997–2001 et al., 2001), we evaluate whether the combined functional hubness and associated microstructural changes could potentially be used as a biomarker to predict post-surgical outcome.

## 2 Results

We analyzed adult participants including 25 patients diagnosed with unilateral pharmaco-resistant temporal lobe epilepsy and 48 healthy controls. All participants underwent an identical 3T MRI protocol which included a 7-minute rs-fMRI run and multishell DWI acquisitions. To map connector hubs for each participant, SPARK was applied to the preprocessed rs-fMRI data, generating subject-level *k*-hubness maps composed of discrete values ranging from 1 to 10, reflecting the number of RSNs contributing to each voxel, as well as atom maps corresponding to canonical RSNs, estimated specifically for each participant. We then identified regions of interest along the cortical surface, unfolded hippocampus, and subcortical structures, demonstrating significant differences in *k-*hubness between TLE and healthy controls, characterizing the distribution of RSNs and calculated the hub disruption index within these regions. Finally, ADC and FA measures derived from DWI were estimated to assess white matter microstructural changes associated with connector hub alterations through group and subject level analyses within the identified regions of interest (see Material and Methods section for further details).

### Global network findings across healthy controls, left- and right-TLE

Using preprocessed resting state fMRI data, connector hub mapping was performed using SPARK to decompose the whole-brain rs-fMRI run of every subject into *N* atom maps, inherently representing *N* canonical resting state networks. For each subject, the atom maps were visually identified and were labeled as either a canonical RSN or as a noisy component (i.e.: cardiac, respiration, scanner signals). Noisy components were subsequently removed, and a final *k-* hubness map was generated for each subject (Lee et al., 2016, 2019). Assessing the total number of atom maps associated with each rs-fMRI run, healthy controls exhibited on average 30.9 ± 2.9 atoms before the identification of noisy atom maps, and 24.5 ± 3.2 atom maps after removing the noisy components, resulting in an average percentage of noisy atoms of 20.6 ± 6.2 %. The left-TLE cohort exhibited on average 28.9 ± 3.8 atom maps before denoising and 22.5 ± 4.4 atom maps after denoising, with an average percentage of noisy atoms of 22.4 ± 9.0 %. Finally, the right-TLE cohort exhibited on average 29.1 ± 4.1 atom maps before denoising and 22.3 ± 4.6 atom maps after removing the noisy components, therefore having a percentage of noisy atoms of 23.8 ± 4.5%.

### Connector hub estimation in healthy controls

The whole-brain pattern of *k-*hubness in healthy controls was computed by averaging the denoised *k-*hubness maps of each control subject. While the individual *k-*hubness maps exhibited more variability in *k-*hubness values, sometimes reaching a value up to *k* = 10, these strong effects were lost when averaging across subjects. By averaging *k-*hubness maps, the goal is to uncover whole-brain patterns of *k-*hubness. In the case of the healthy cohort in the current study, the maximum value of the average *k-*hubness (*<k>*) was found to be three. The highest *<k>* values across the neocortex were found in the medial prefrontal cortex, posterior cortex/precuneus, and inferior parietal lobe. The hippocampal-amygdala transition area yielded the highest *k-*hubness values along the unfolded hippocampal surface. Subcortical structures demonstrate overall lower *<k>*, with the bilateral putamen demonstrating the highest values (Figure 1A).

**Figure 1:**
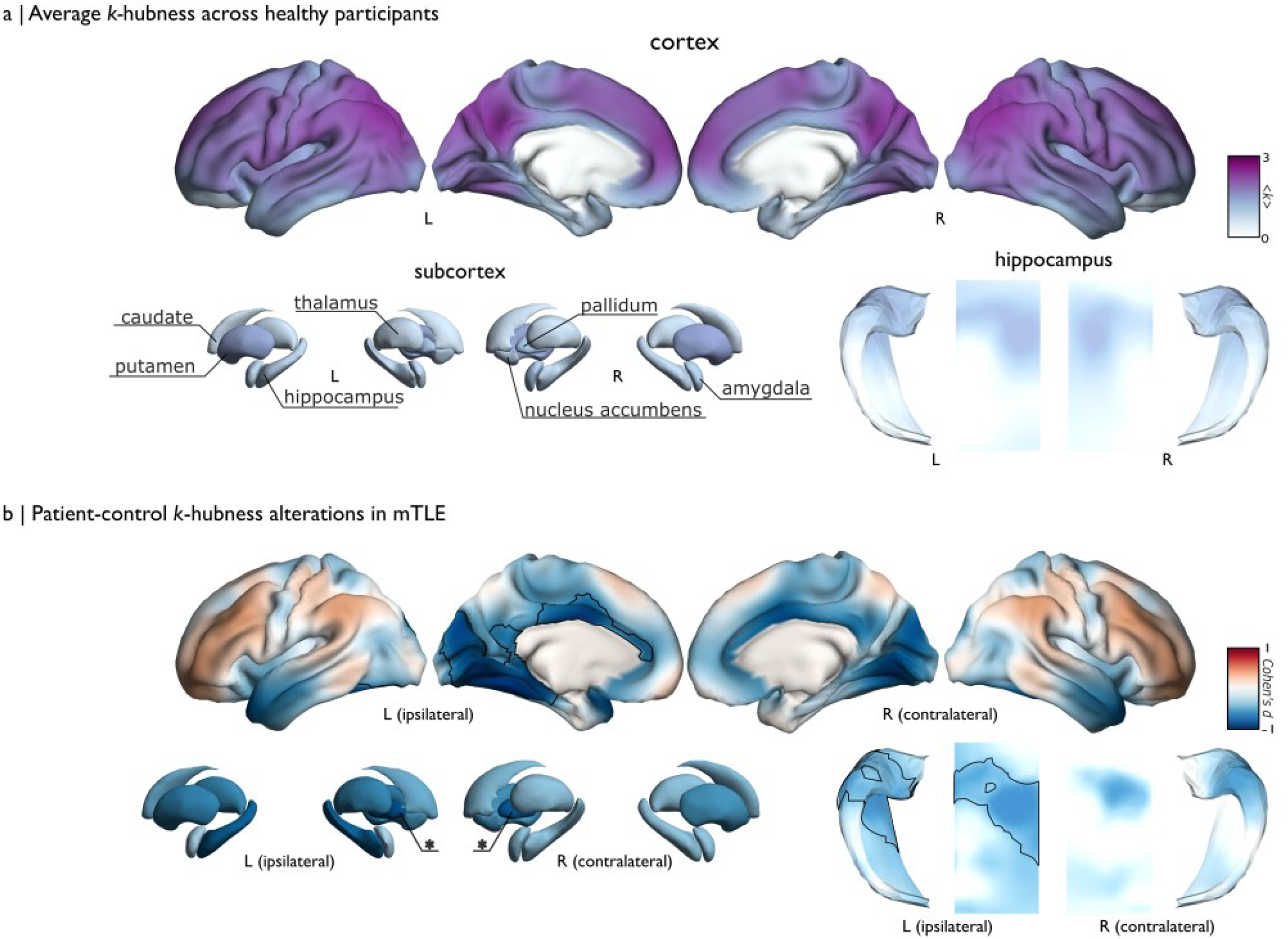
Distribution of *k*-hubness. A | Average *k-*hubness (<*k*>) across healthy participants in the cortex, subcortical structures and the unfolded hippocampus. Color bar between 0, meaning unreliable/no *k-*hubness signal and 3 which is the maximum <*k*> value in the healthy cohort. B| Patients versus controls *k-*hubness alterations in TLE. Outlined in black on the cortex and unfolded hippocampus is the cluster 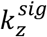 exhibiting significant differences using a two-sided t-test (p_FWE_ < 0.05) and starred in the subcortex (*p*_*FDR*_ *< 0*.*05*). Colormap shows Cohen’s *d* at each vertex, which was computed using 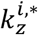 between epilepsy and control subjects ranging between −1 to 1.

### Patients versus controls *k-*hubness alterations in temporal lobe epilepsy

To compare *k-*hubness between groups, we first Z-scored the individual *k-*hubness maps relative to controls, sorting ipsilateral/contralateral to the epilepsy focus, and lateralizing all epilepsy foci to the left hemisphere. Correcting for age and sex using a fixed effect linear model, we obtained vertex-level Z*-*score *k-*hubness maps for each subject 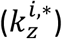. Using these adjusted maps, we evaluated group differences 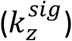 while correcting for multiple comparisons, and calculated Cohen’s *d* at each vertex. Figure 1B illustrates these effect sizes at each vertex, with the significant cluster 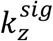 outlined in black. Statistical significance was assessed using *t*-tests thresholded at *p* < 0.05. To correct for multiple comparisons, we applied Random Field Theory based using Family-Wise Error (FWE) correction for surface-based analyses and False Discovery Rate (FDR) correction for volumetric analyses. Patients with TLE demonstrated significant widespread decreases of *k-* hubness in the medial occipital region (cuneus, lingual gyrus), the mesial temporal regions (parahippocampal gyrus, fusiform gyrus, and inferior temporal gyrus), the cingulate gyrus and the precuneus (*p*_*FWE*_ < 0.05, *mean* ± *SD* Cohen′s *d* = −0.16 ± 0.26). Along the hippocampal surface, we found significant decreases in the ipsilateral hippocampal head (*p*_*FDR*_ < 0.05, mean ± SD Cohen’s *d*_*ipsi*_ = −0.33 ± 0.21). Subcortical analysis uncovered significant hubness decreases in the bilateral pallidum (*p*_*FDR*_ < 0.05, Cohen′s *d*_*ipsi*_ = −0.70, Cohen′s *d*_*contra*_ = −0.67).

### Resting state networks involved with regions associated with significant hub changes

Benefiting from the unique ability of SPARK to identify specific RSNs associated to each hub, we generated a dataset specific RSN atlas (SPARK_RSN_) from healthy control data. To build SPARK_RSN_, each atom map across all subjects was manually assigned to a canonical resting state network. For each functional network defined in the SPARK_RSN_ atlas, we averaged all instances across healthy subjects and thresholded the resulting map to keep its most spatially reliable regions. For each element in SPARK_RSN_, we computed the area of spatial overlap between the network and significant clusters 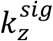. In total, we identified fifteen reoccurring networks across all participants. In Figure 2, we visualize the networks contributing to more than 2% of the total area of the significant clusters. In 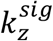 along the cortical surface, a total of seven networks were identified: the visual medial (overlap = 60.47%), visual occipital (overlap = 33.59%), visual lateral (overlap = 33.48%), precuneus (overlap = 10.84%), default mode (overlap = 8.46%), salience (overlap = 8.01%) and temporal (overlap = 2.92%) networks (Figure 2A). Along the unfolded hippocampal surface, six networks were found to contribute to the hippocampal cluster: hippocampus (overlap = 100%), temporal (overlap = 97.30%), default mode (overlap = 64.83%), subcortical (overlap = 33.29%), precuneus (overlap = 3.33%), and auditory (overlap = 2.98%) networks (Figure 2B). The bilateral pallidum was found to only contribute to the subcortical network (overlap_ipsi_ = 98.11%, overlap_contra_ = 96.93%) (Figure 2C).

**Figure 2:**
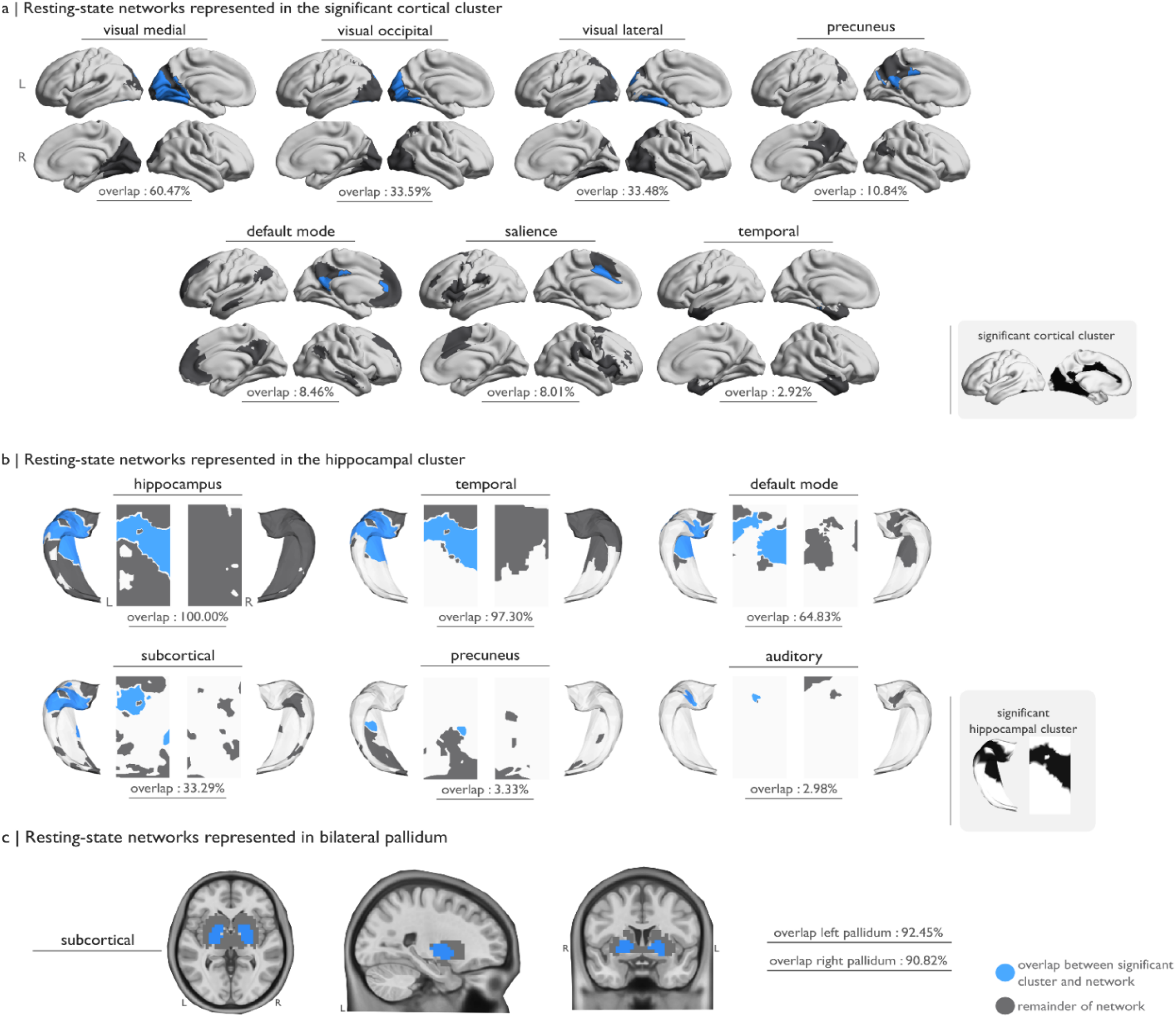
Networks from SPARK_RSN_ which are identified to be overlapping with the significant clusters 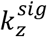 in the neocortex, along the unfolded hippocampus and for subcortical structures. Visualized networks all exhibited more that 2% overlap between 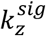 and the network in SPARK_RSN_. In the visualization, blue is a vertex where both 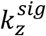 and the SPARK_RSN_ are overlapping and grey is the remainder of the corresponding SPARK_RSN_ network. A | The SPARK_RSN_ networks associated with the neocortical 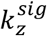 region. Networks include visual medial, visual occipital, visual lateral, precuneus, default mode, salience and temporal SPARK_RSN_ networks. B| The SPARK_RSN_ networks associated with the unfolded hippocampal 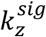 cluster, including hippocampal, temporal default mode, subcortical, precuneus, and auditory networks. C | The subcortical SPARK_RSN_ network was the only one to be associated with the subcortical 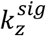.

### Hub disruption index

We adapted the hub disruption index introduced in (Achard et al., 2012; Lee et al., 2018) to be applied alongside the age-and-sex adjusted data, 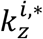. We computed a linear regression model between the normalized mean of the hemispheric average of controls (x-axis) and the Cohen’s *d* between epilepsy and controls (y-axis). The slope of this regression models how hubness changes relative to the distance from the average of controls. Relative to healthy controls, a negative slope represents hubs in patients are losing hubness, whereas a positive slope indicates hubs in patients are gaining hubness. Furthermore, a positive intercept represents hubs in patients are initially demonstrating higher hub values compared to the average of controls, and a negative intercept indicates that hubs in patients are initially demonstrating lower hub values compared to the average of controls. We computed this regression model on the three ipsilateral 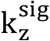 and their homologous structures on the neocortex (Figure 3A), unfolded hippocampus (Figure 3B) and subcortical volumes (Figure 3C). To account for spatial autocorrelation when testing the significance of the hub disruption index, we performed spin tests on the neocortical and unfolded hippocampus data, and applied Moran’s Randomization on the subcortical volumes (*p* < 0.05). All regions demonstrated negative slopes suggesting an overall loss of connections in TLE patients. The slopes in contralateral homologous regions in the neocortex and hippocampus exhibited much more negative trends 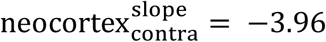, p < 0.05, 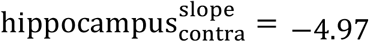, p < 0.001) than their ipsilateral counterparts 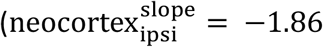, p < 0.05, 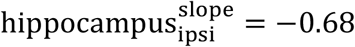, no significance), implying a wider range of Cohen’s *d* values in the contralateral regions. These negative slopes can be interpreted as hub regions in controls loosing their hubness in patients. In the neocortex and unfolded hippocampus, ipsilaterally, 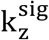 demonstrated negative intercepts 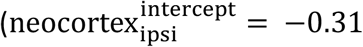, no significance, 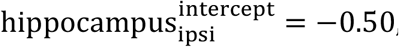, no significance = −0.50), while their contralateral homologous regions demonstrated positive intercepts 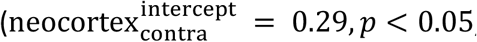, *p* < 0.05, 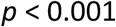, *p* < 0.001). These results are suggesting contralateral initial hub emergence in regions that are considered non-hubs in healthy controls through additional connections being made before rapidly losing connections due to their negative slope. The bilateral pallidum ROIs exhibited intercepts around zero and similar negative slopes bilaterally.

**Figure 3:**
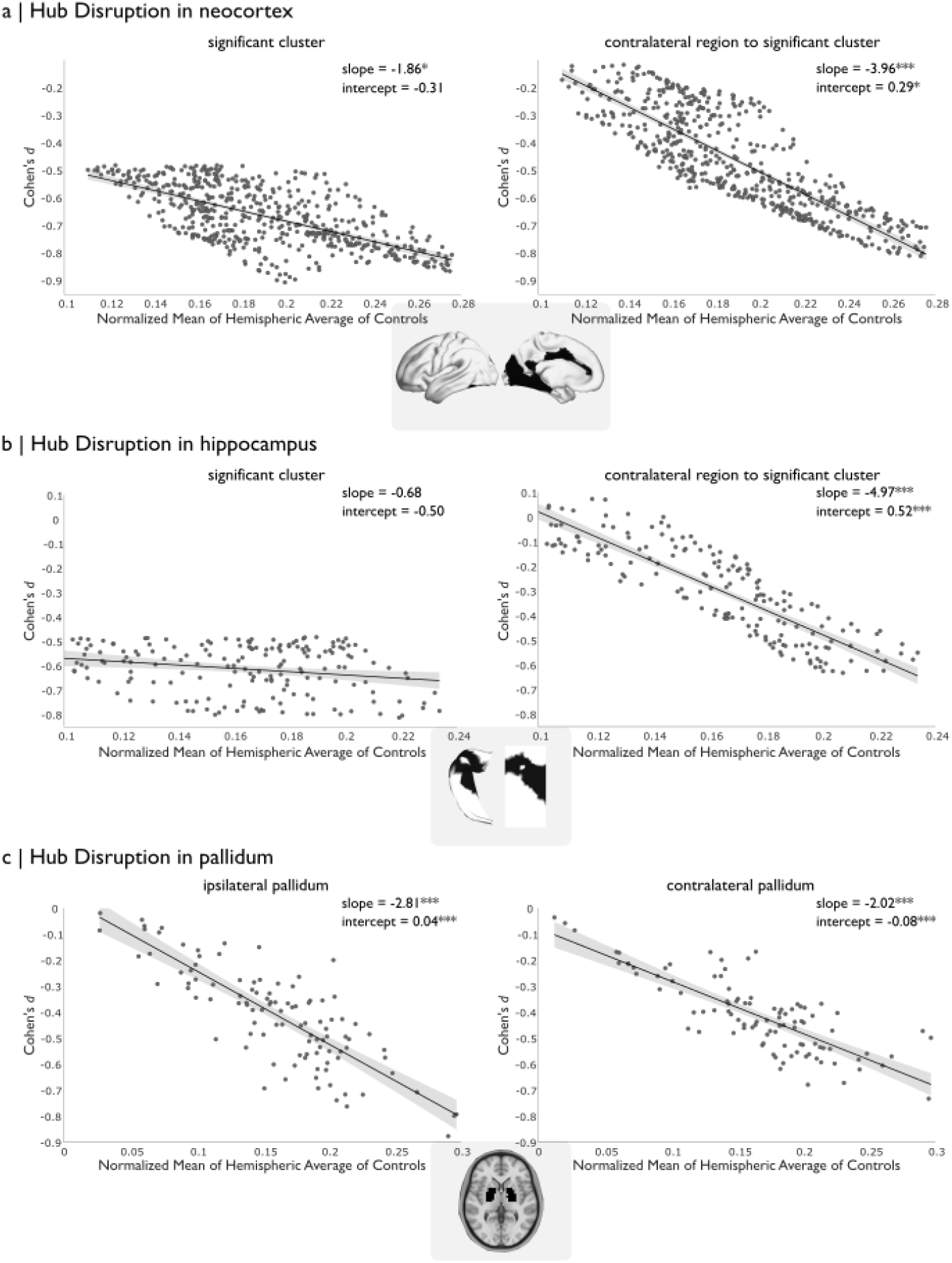
Hub disruption index. X-axis is the normalized mean of the hemispheric average of controls 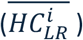 and the y-axis is the Cohen’s *d* comparing patients versus controls for each vertex within the region of interest. The slope of the plot represents HDI, quantified as the rate of change of *k-*hubness, and the intercept represents the pattern of the *k-*hubness value when there is zero variation within the healthy control group (baseline level of healthy controls, Z = 0), quantifying whether the region experiences initial disruption or emergence. A| The hub disruption index within the 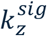 of the neocortex and its contralateral homologous region. B| The hub disruption index within the 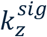 of the unfolded hippocampus (ipsilateral and contralateral). C| The hub disruption index within the subcortex’s 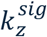 regions. Significance testing was performed using spin tests. \**p < 0*.*05, ** p < 0*.*01, ***p < 0*.*001*

### Patients versus controls microstructural white matter changes in temporal lobe epilepsy

In Figure 4, we identified microstructural white matter changes in TLE patients versus controls using apparent diffusion coefficient (ADC) and fractional anisotropy (FA) extracted from diffusion weighted imaging. Following the same procedure we applied for *k-*hubness, Figure 4A shows the significant univariate microstructural changes after regressing out the effect of age and sex, where 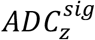 (top) and 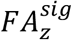 (bottom) are outlined in black overlayed on the Cohen’s *d* maps. For ADC, we found widespread significant increases in the ipsilateral ventromedial prefrontal cortex, temporal lobe, hippocampus and amygdala. FA demonstrated more localized significant decreases limited to the temporal pole, medial occipitotemporal gyrus, and orbitofrontal cortex. We then recomputed 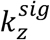 after regressing out MRI-derived microstructural white matter effects at a vertex level in Figure 4B (top). In this recomputed cluster comparing *k-*hubness in TLE patients versus controls, we found that 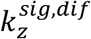 reduced in size in the anterior cingulate gyrus, whereas 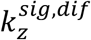 for the unfolded hippocampus and the bilateral pallidum no longer exhibited any significant differences between groups. We also investigated the patient-specific *k-*hubness profiles, where we compared the average 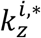 before and after the correction of microstructural white matter in the ipsilateral and contralateral homologous regions of 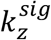 along the cortical surface, the whole cortex, the unfolded hippocampus, the bilateral pallidum, and the entire subcortex. In Figure 4B (bottom), we show that ipsilateral ROIs closer and/or involved with the mesial temporal lobe, demonstrated weaker Cohen’s *d* values after correcting for microstructural white matter effects of *k-*hubness. Particularly, the ipsilateral hippocampus (Cohen’s *d*_uncorrected_ = −0.45, Cohen’s *d*_corrected_ = −0.33) demonstrated the strongest effect. We found no significant effect between the groups when comparing before and after correction of diffusion metrics across all ROIs.

**Figure 4:**
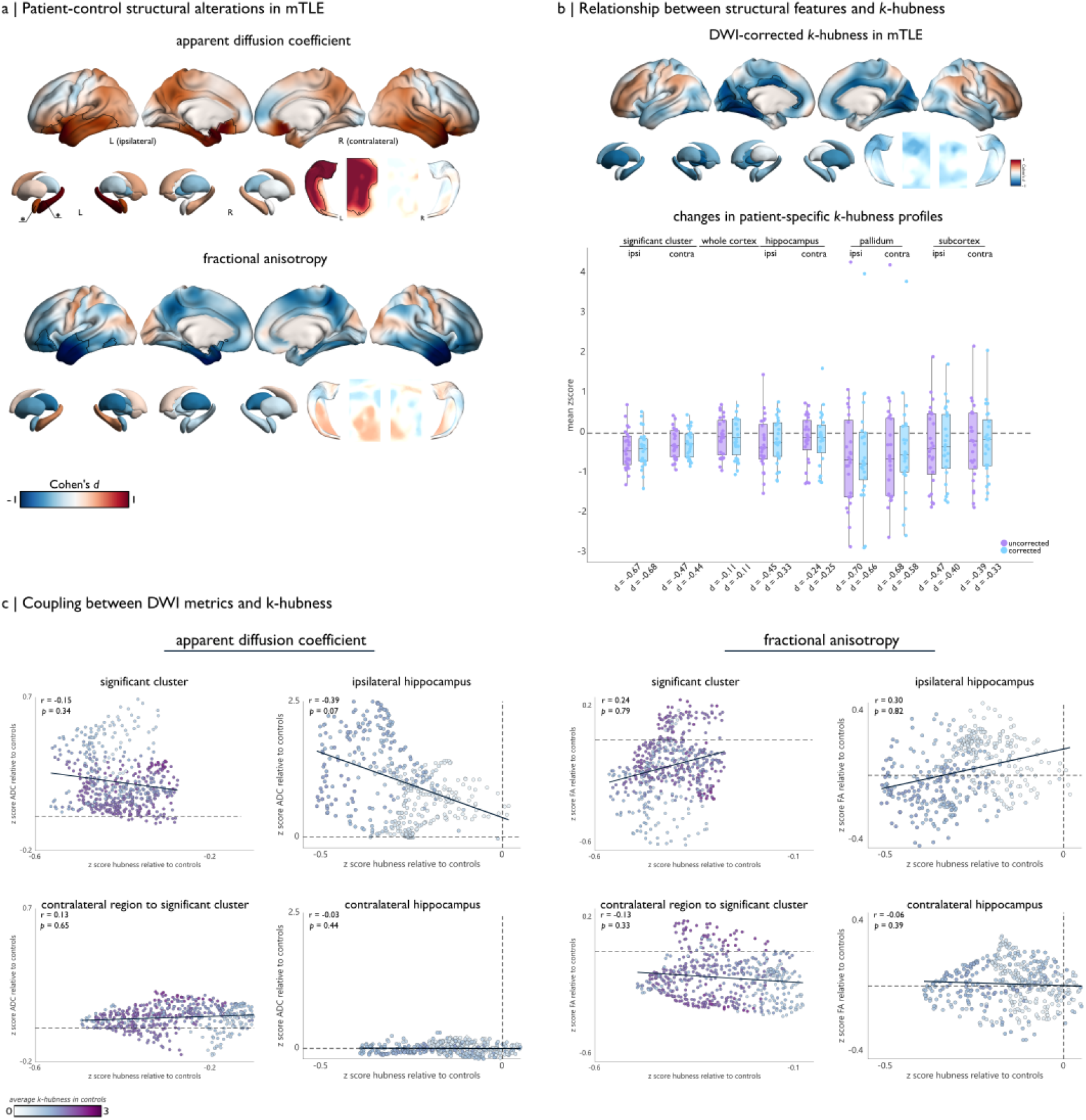
The relationship between *k-*hubness and white matter microstructure. A | Univariate microstructural white matter changes in TLE relative to healthy controls. 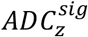 (top) and 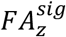 (bottom) are outlined in black on the neocortex and unfolded hippocampus (*p* < 0.05, FWE corrected), and starred in subcortex (*p* < 0.05, FDR corrected). Maps are displaying Cohen’s *d* between epilepsy and control groups using 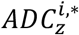 and 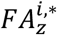 maps which were corrected for age and sex. Color bar ranging between −1 to 1. B | The relationship between *k-*hubness and white matter microstructure. Top: We recomputed 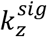 after also correcting for ADC and FA diffusion metrics in the neocortex, subcortex and unfolded hippocampus. Bottom: Changes in patient-specific *k-*hubness profiles where each point represents the average of the vertices found in 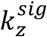.Purple represents the original 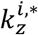 data, and blue represents the ADC and FA corrected 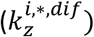 data. The average Cohen’s *d* within 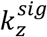 before and after correction is displayed underneath the boxplots. C | Coupling between microstructural metrics and *k-*hubness within the neocortical 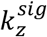and the contralateral homologous region and the bilateral hippocampi. Coupling between 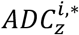 and 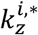 is on the left panel and coupling between 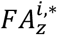 and 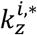 coupling on the right of the panel. Color bar is < *k* > in healthy controls from Figure 1A, ranging between 0 and 3.

### Correlation between *k-*hubness and microstructural white matter metrics in TLE

We estimated the Pearson’s correlation (*r*) between the 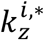 of patients and the corresponding 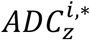 and 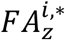 maps in the 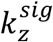 of the neocortex and unfolded hippocampus along with their corresponding contralateral homologous regions. Within ipsilateral significant regions, our results suggested a negative correlation between ADC and *k-*hubness (decrease *k-*hubness associated with an increase of ADC) and a positive correlation between FA and *k-*hubness, however these trends did not reach significance. Contralateral homologous regions did not exhibit the same patterns, where the z-scored diffusion metrics collapsed around zero. The strongest coupling relationship we observed was between ADC and *k-*hubness in the ipsilateral hippocampus (*r* = −0.39, *p*=0.07), where the vertices representing the strongest hubs in controls were found to exhibit the strongest alterations in both metrics exhibiting decreased *k-*hubness and increased ADC (see Figure 4C).

### Applying the model to seizure free patients

In Figure 5A, we repeated our analysis in a small subset of patients (n=9) who underwent surgical intervention and have been seizure free after one year follow-up (Engel IA). When considering this small subset of patients, we estimated a significant cluster 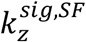 between patients and controls that was concentrated around the mesial temporal and medial visual structures, with a minor contribution in the middle-cingulum after regressing out the effect of age and sex. When extending to the unfolded hippocampal surface, our results exhibited a much more extended region in the hippocampal body contributing to significant decreases in *k-*hubness. The subcortical structures exhibited much more widespread significant decreases in *k-*hubness, extending through the bilateral accumbens, pallidum, putamen, thalamus, and the ipsilateral hippocampus. Analysis of structural features (ADC, FA) demonstrated significant differences compared to controls in the ipsilateral temporal lobe. All regions were estimated at a threshold of *p < 0*.*05* and were corrected for multiple comparisons.

**Figure 5:**
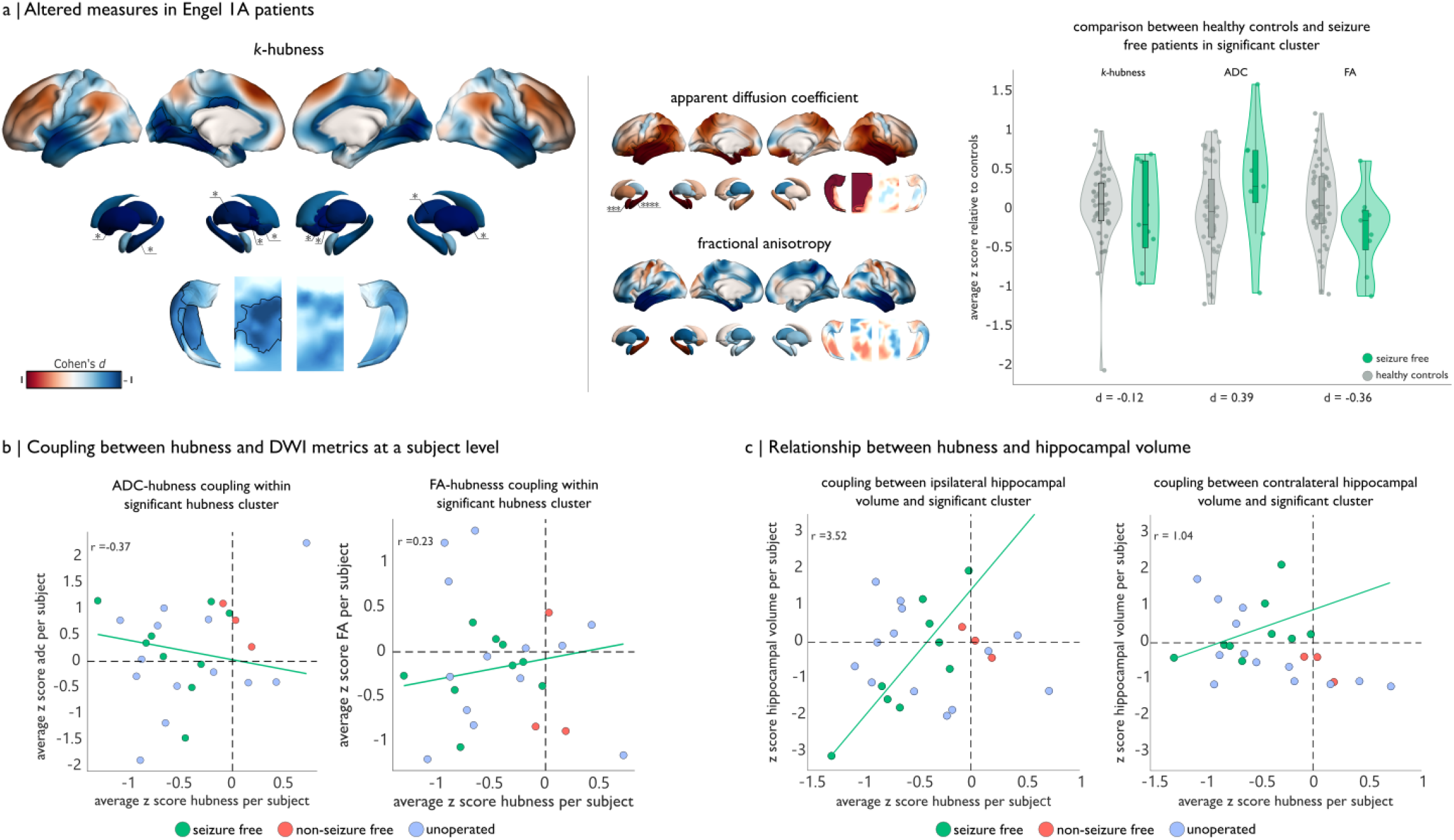
The relationship between seizure freedom with *k-*hubness, ADC, FA and hippocampal volume. A| The significant clusters 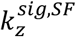 (left), 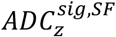 (middle-top), and 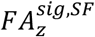 (middle-bottom) outlined in black overlayed on Cohen’s *d* maps when rerunning our analysis on a subgroup of 9 seizure-free patients in the neocortex, unfolded hippocampal surface. (*p* < 0.05, FWE corrected) and subcortex (*p* < 0.05, FDR corrected). Color bar ranges from −1 to 1. On the right of the panel, violin plots display subject-level average of 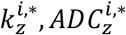, and 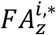 within the 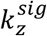 from Figure 1B. Seizure free patients are in green and healthy controls are in grey. B| Subject-level coupling between *k-*hubness and microstructural metrics in the 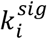 from Figure 1B. Seizure free patients are in green, non-seizure free patients are in red, and unoperated patients are in blue. Green line is the linear regression line estimated when considering only the seizure-free patients. C| Relationship between *k-*hubness and hippocampal volume, following the same color scheme and regression when considering seizure free patients only from panel B.

We also reproduced the functional-structural relationship at the individual level within the cortical 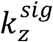 from our primary analysis (Figure 1B), where we averaged the 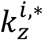 values for each subject. In Figure 5B, we are showing the profiles for all patients, featuring the seizure-free patients are in green, non-seizure free patients in red, and unoperated patients in blue. When assessing the correspondence between ADC and *k-*hubness, the green trendline which was fit only on the seizure-free patients, follows the expected pattern that we observed at a group level in Figure 4C. We find 6 out of the 9 seizure-free patients closely following the trend, and all nine subjects demonstrated reduced *k-*hubness relative to controls. Also including the 3 non-seizure free subjects in the analysis, two of the three non-seizure free patients demonstrated an increase of *k-*hubness relative to controls, while still demonstrating the expected changes in ADC and FA.

### Relationship between *k-*hubness and hippocampal volume

As an additional structural biomarker, we also examined the relationship between hippocampal volume and *k-*hubness at the individual patient level in the 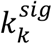 from Figure 1B. We found a direct relationship between hippocampal volume and *k-*hubness, where a decrease in hippocampal volume relative to controls corresponds to a decrease in *k-*hubness. Contralaterally, the relationship was present but not as pronounced, with less variation in hippocampal volume across all patients.

## 3 Discussion

In the present study, we applied *SParisty-based Analysis of Reliable k-*hubness (SPARK), a functional connectivity framework for mapping reliable connector hubs (Lee et al., 2016, 2018, 2022, 2026), to characterize alterations in *k-*hubness in patients with unilateral temporal lobe epilepsy relative to healthy controls. We identified widespread reductions of cortical *k-*hubness across the mesial occipital, temporal, and cingulate structures, accompanied by decreases throughout the ipsilateral hippocampus and in the bilateral pallidum of the subcortex, therefore reproducing and extending our earlier findings applying SPARK on a more limited mTLE database (Lee et al., 2018). To better characterize the functional organization of these changes, we leveraged the uniqueness of SPARK to uncover each individual subject’s resting state networks to generate a dataset-specific atlas of resting state networks and associate it with regions exhibiting significant loss in connector hubs. Here, we revealed that altered hub regions were predominantly associated with multiple visual networks, while the default mode network was uncovered as a major contributor across the neocortex and unfolded hippocampus. We further demonstrated that changes in hippocampal and subcortical *k-*hubness were more strongly associated with underlying white matter microstructure than those observed along the cortical surface. Finally, an analysis of a subgroup of seizure-free patients revealed more localized differences of *k-* hubness along the cortex but more widespread reductions in hippocampal and subcortical *k-* hubness.

### The role of resting state networks

In this work, we are reporting how significant connector hub differences were found to spatially overlap with various RSNs spread across the neocortical and unfolded hippocampal surfaces. Using our hub mapping method SPARK, we have the unique capability of understanding which RSNs are associated with each hub, allowing us to generate a dataset specific atlas (SPARK_RSN_) of networks. First of all, using the SPARK data driven exploration technique, we retrieved our SPARK_RSN_ atlas by identifying several canonical RSNs reported in the literature in healthy controls (Heine et al., 2012; Di and Biswal, 2014; Lv et al., 2018). Our results are suggesting that the three most impacted RSNs contributing to areas of significant hub changes in the neocortex were the three spatially distinct visual networks. Previous work has shown how occipital regions have demonstrated increased within-network connectivity (Zhang et al., 2009) and decreased in inter-network connectivity (Dumlu et al., 2020) in temporal lobe epilepsy, supporting our findings. This can possibly suggest a local-hub reorganization, where connector hubs are losing their long-range connections and are instead forming more local, provincial hubs. Moreover, our analysis revealed that significant clusters demonstrating *k-*hubness decreases in TLE patients overlapped with both anterior and posterior default mode network regions across neocortical and hippocampal surfaces. Our results further support prior work demonstrating decreased connectivity between these default mode regions (Haneef et al., 2012; Cataldi et al., 2013), suggesting a disruption or loss of connector hubs in TLE. Additionally, the default mode has been shown to exhibit decreased functional connectivity with the hippocampus and limbic structures (Pittau et al., 2012; de Campos et al., 2016), in agreement with our observations and reproducing our previous results suggesting hub disruption in the default mode network (Lee et al., 2018). Furthermore, SPARK uncovered a precuneus network, spatially unique from the posterior default mode which was found to exhibit an overlap with regions of significant decreases in *k-*hubness in both the neocortex and unfolded hippocampal surfaces. Our team has previously shown the precuneus to be a high-hub region in healthy controls (Lee et al., 2016), in a study assessing the test/retest reliability of SPARK, a result that we are reproducing here using SPARK on a more extended database. The precuneus has shown to demonstrate increased within-network connectivity (Haneef et al., 2012; Zanão et al., 2021; Zhou et al., 2021; Huang et al., 2022), and reduced long-range connectivity of connector hubs (Liao et al., 2011; McCormick et al., 2013), consistent with our findings. Finally, the dorsal anterior cingulate portion of the salience network also exhibited a significant decrease in *k-*hubness, a region which has previously demonstrated reduced inter-network connectivity and plays a mediating role between the default mode and the executive networks (Zhou et al., 2021; González et al., 2022). Our findings of multi-network interactions with regions exhibiting significant decreases of *k-*hubness in TLE reinforce the viewpoint of how TLE is a network-level disorder. The combination of the reduced inter-network connectivity we are presenting in this work in combination with reports of increased within-network connectivity throughout the literature supports a model of hub reorganization where widespread disruptions in connector hubs lead to more locally constrained network architectures. In epilepsy, this phenomenon has been previously described as a localized regular network which has also been found in across multimodal analyses (Chiang and Haneef, 2014). Moreover, the presence of key networks such as the default mode and precuneus within pathological regions may further exacerbate inter-network disconnection due to their central, overlapping roles across multiple functional systems.

### Hub disruption index can be used to quantify the rate of how hubs change

We used the hub disruption index to quantify the rate of hubness changes within the significant cluster and their contralateral homologous region in our analysis. This metric was originally introduced by (Achard et al., 2012), who demonstrated that regions acting as hubs in healthy controls lost connections in comatose patients. This framework was later extended by (Lee et al., 2018), who introduced the concept of hub emergence, showing that non-hub regions in controls gained connectivity in patients with right mTLE. Building upon these previous studies, we adapted the HDI metric to fit within our proposed framework in Figure 3, by applying a linear regression between the normalized mean of the hemispheric average of controls (x-axis) and the Cohen’s *d* between epilepsy and controls (y-axis). Consistent with prior work, we observed evidence of widespread hub disruption, where our clusters are exhibiting progressive losses of connectivity relative to controls. However interestingly, contralateral ROIs were associated with positive intercepts, suggesting that regions with relatively low hubness in controls may initially gain connectivity before undergoing disruption. This may be indicative of a compensatory response to the ipsilateral pathology, whereby an initial compensatory effect is to develop hubs to support network function, as proposed by Bettus et al., 2009; an effect also associated with good cognitive surgical outcome (Doucet et al., 2015). Notably, the same contralateral regions also demonstrated rapid hub disruption, indicating that although compensatory hub formation may occur, it is insufficient to prevent an overall decline in connectivity relative to controls. In the hub disruption plots, the contralateral slopes were indeed steeper than the ipsilateral ones. This most likely reflects greater variation in Cohen’s *d* values across contralateral regions, whereas ipsilateral regions were selected based on significant hub disruption. The fact that ipsilateral regions have a slope implies that even though the hubs have already undergone a significant disruption, they are still capable of losing even more connections. The emergence of connectivity in contralateral regions has received comparatively little attention outside of the work by (Lee et al., 2018). Our findings therefore extend previous HDI studies in epilepsy, which have consistently reported negative whole-brain HDI values across rs-fMRI (Ridley et al., 2015; Song et al., 2015) and across frequency bands in the context of high-density EEG source analysis in generalized epilepsy (Alves et al., 2024).

### White matter microstructure contributes to hippocampal and subcortical *k-*hubness

We also evaluated the impact of the underlying white matter microstructure on our functional measure of *k-*hubness. Our results reflecting the univariate changes in white matter microstructure unveiled concordant significant changes in TLE relative to controls consistent with previous literature (Otte et al., 2012; Bernhardt et al., 2016; Hatton et al., 2020; Ngo et al., 2024; Chen et al., 2025); mainly consisting in significant increases of ADC and decreases of FA localized to the ipsilateral temporal structures, in comparison to the more widespread changes of *k-* hubness. To explore how long-range connections in these regions depends on the underlying white matter microstructure, we recomputed our surface based linear model by regressing out FA and ADC before re-estimating the group comparison. While we found minimal changes in the regions showing significant hub disruption in the cortex, both the unfolded hippocampal and pallidum regions were no longer associated with atypical hub organization. These findings reflects the biological dependency of these regions, as hippocampal functional connectivity has been shown to be closely associated with the integrity of the white matter pathways (Kehoe et al., 2015). Furthermore, our significant region along the hippocampal surface also exhibited some overlap with the default mode network, which is a network who’s functional connections has been shown to be coupled with white matter tract integrity. (Teipel et al., 2010). Additionally, the structure-function coupling effects in both the neocortical and unfolded hippocampal clusters showing significant hub disruption revealed ipsilateral coupling between the functional *k-*hubness changes and white matter microstructural alterations, whereas contralateral homologous regions exhibited some functional changes in the absence of detectable white matter abnormalities. These effects were primarily localized to regions that correspond to high-hub areas in healthy controls, suggesting preferential vulnerability of network hubs to structure-function reorganization in TLE. This is consistent with evidence that network hubs are disproportionally implicated across a broad range of brain disorders, with structural abnormalities preferentially concentrated in highly connected rich-club regions (Crossley et al., 2014). In TLE specifically, previous work has shown that rich-club functional hubs demonstrate overlap with structural changes in the cortex (Liu et al., 2016; Larivière et al., 2020), and the hippocampus (Chiang et al., 2015) supporting the current findings of connector hub lead coupling.

### An exploratory look at hubness and surgical outcome in TLE

Lastly, we performed an exploratory trifold analysis evaluating the relationship between *k-* hubness, white matter microstructure and postoperative seizure freedom. To do so, we recomputed our model where we focused on a limited subgroup of nine patients who have achieved seizure freedom after surgery. We found that subcortical structures including the bilateral pallidum, putamen and thalami, as well as the ipsilateral hippocampus showed more widespread decreases in *k-*hubness relative to our cohort of control participants. Our results further revealed more localized cortical regions of significance, including medial visual cortex and middle cingulum, alongside mesial temporal structures. While limited by a small cohort size, these findings are highly encouraging and as they mimic key regions of an epilepsy propagation network previously identified as predictors of favorable surgical outcomes in functional connectivity studies (Morgan et al., 2017, 2019). Additionally, pre-surgical thalamic hubness has been established as a predictor of post-surgical seizure freedom. When compared to controls, patients with higher pre-surgical hubness exhibit poorer outcomes, whereas those with lower hubness achieve positive ones (He et al., 2017). Our group level results are consistent with and further support these findings, where we report significantly reduced bilateral thalamic *k-*hubness in seizure-free patients. Notably, many of the regions associated with seizure freedom overlapped with those identified in our primary group analysis, suggesting that *k-*hubness may capture network reorganization relevant to clinical outcome. Future work examining the differences between seizure-free and non-seizure free patients, particularly in regions that do not overlap with the broader TLE-related alterations will be important for determining whether *k-*hubness captures outcome-specific patterns of network reorganization. Furthermore, the widespread involvement of subcortical and cortical regions beyond the mesial temporal lobe supports the growing recognition of TLE being a disorder of brain networks rather than a condition confined to the mesial temporal lobe (Spencer, 2002; DeSalvo et al., 2014; Bernhardt et al., 2015; Bartolomei et al., 2017; Whelan et al., 2018). While limited, our individual level coupling analysis demonstrated consistent decreases in *k-*hubness in seizure-free subjects while the non-seizure free subjects showed similar or increased *k-*hubness relative to healthy controls. This pattern is consistent our previous study reporting similar alterations in hubness within the epileptogenic zone (Aydin et al., 2020). Furthermore, Karpychev et al. (2026a) demonstrated that long-term surgical outcomes depend not only on the accurate localization of the epileptogenic zone, but also on its integration within the broader brain network. Specifically, they showed that greater cross-network connectivity within the epileptogenic zone is associated with a higher likelihood of seizure recurrence. Interestingly, our functional connectivity result contrasted with our individualized white matter microstructural findings. While seizure-free patients did not uniformly exhibit the expected microstructural alterations (i.e., increased ADC and decreased FA relative to controls), all the non-seizure free patients did. This discrepancy suggests that functional network alterations and white matter microstructure may capture distinct, yet complementary aspects of the disease pathology. Whereas diffusion abnormalities likely reflect underlying tissue damage (Rodríguez-Cruces and Concha, 2015), *k-*hubness may more directly characterize the large-scale functional architecture. Notably, within our small cohort, individualized *k-*hubness profiles better distinguished seizure-free from non-seizure free patients than diffusion-derived microstructural measures, suggesting that functional network organization may provide credible information beyond conventional structural biomarkers. More broadly, our results suggest that connector hub mapping provides clinically relevant information that complements structural imaging by quantifying the integrity of distributed functional networks. While requiring validation in larger prospective cohorts, individualized assessments of *k-*hubness may ultimately contribute to improved presurgical stratification and prediction of postoperative seizure outcomes.

### Limitations

Several limitations arise when interpreting our findings. First, although we sought to maximize cohort homogeneity by restricting our analyses to patients with unilateral pharmaco-resistant TLE, important sources of clinical variability remain unavoidable. Particularly, our cohort was not controlled for the number and type of antiseizure medications, and the underlying pathology was not uniform across patients. Similarly, the subgroup of seizure free patients underwent different surgical procedures, which may have influenced postoperative network organization and contributed additional variability to our outcome analysis. Moreover, while previous work from our group has demonstrated distinct patterns of connector hub disruption in left and right lateralized mesial temporal lobe epilepsy (Lee et al., 2018), we elected to lateralize all patients to a common hemisphere rather than performing separate analyses. Given the relatively small cohort size (15 left and 10 right lateralized patients), this approach increases statistical power, enabling a more robust investigation of the relationships between functional hub organization and white matter microstructure. Future studies with larger cohorts should revisit lateralization differences to determine whether the functional-structural relationships identified here are influenced by seizure laterality. Methodologically, this study focused exclusively on alterations in connector hubs. Throughout our analyses, we observed patterns suggestive of a transition from connector to provincial hub organization. However, provincial hubs were not explicitly quantified. Future work incorporating complementary measures of provincial hub architecture would allow this hypothesis to be directly tested and provide a more comprehensive characterization of network reorganization in temporal lobe epilepsy. Finally, this was a single site cohort and included a relatively modest sample size, particularly for the postoperative seizure analysis. Although many of our findings are consistent with previous studies, replication in a larger, independent, multi-site cohorts would be essential to establish the robustness, generalizability, and potential clinical utility of connector hub mapping as a biomarker in epilepsy.

## 4 Materials and Methods

### Participants

Twenty-five (25) adult patients diagnosed with pharmaco-resistant unilateral temporal lobe epilepsy (Age: 40.28 ± 12.07 yrs, 21 right-handed, 14 females), and forty-eight (48) healthy control participants (Age: 34.58 ± 10.66 yrs, 45 right-handed, 24 females) were recruited between 2018 and 2024 at the Montreal Neurological Institute and Hospital. Following our research protocol, all subjects underwent identical 3T MRI brain acquisitions, with parameters similar to the MICA-MICs dataset reported in Royer et al., 2022b. Patients were included based on the availability of both rs-fMRI and DWI scans, as well as exhibiting an average framewise displacement (FD) < 0.05 mm during their rs-fMRI run. In this cohort, 15 patients had left-TLE and 10 patients had right-TLE. Control participants were free of any neurological or psychiatric conditions. A TLE diagnosis for each patient was determined after staying at the epilepsy monitoring unit, where they underwent comprehensive evaluation that included clinical history, neurological examination, review of medical records, prolonged video-EEG telemetry recordings, clinical-neuroimaging, and neuropsychological evaluation. The diagnosis was determined according to the International League Against Epilepsy criteria. All patient MRI data was acquired prior to surgery. A subset of patients (n = 12) underwent surgical intervention after their initial workup, where 9 were diagnosed as seizure free (Engel IA) and 3 were non-seizure free one year after surgery.

### MRI Acquisition

MRI data in patients and healthy controls was acquired on a 3T Siemens Magnetom Prisma-Fit equipped with a 64-channels head coil. For all subjects, the acquisition protocol included: (i) high-resolution 3D T1-weighted (T1w) scans using a magnetization-prepared rapid gradient-echo sequence (MPRAGE) (repetition time [TR] = 2,300 ms, echo time [TE] = 3.14 ms, flip angle [FA] = 9°, matrix=320 × 320, voxel size = 0.8 × 0.8 × 0.8 mm^3^, field of view [FOV] = 256 × 256 mm^2^, 224 slices); (ii) resting-state fMRI scans using a multiband accelerated 2D EPI sequence (TR = 600 ms, TE = 30 ms, FA = 52°, FOV = 240 × 240 mm^2^, voxel size = 3 × 3 × 3 mm^3^, matrix size = 80 × 80, multiband factor = 6.48 slices, 700 volumes); and (iii) diffusion weighted MRI using a 2D EPI sequence (TR = 3500 ms, TE = 64.4 ms, FA = 90°, FOV = 224 × 224 mm^2^, voxel size = 1.6 × 1.6 ×1.6 mm^3^, 3 b0 images, b-values = 300/700/2000 s/mm^2^ with 10/40/90 diffusion directions).

### MRI Preprocessing

Multimodal MRI data processing was performed using micapipe_v0.2.0 (Cruces et al., 2022). T1w images, were reoriented to LPI orientation (left-right, posterior-anterior, inferior-superior), underwent intensity non-uniformity correction (N4) and intensity rescaling de-obliqued. Subcortical structures were segmented using FSL FIRST (Patenaude et al., 2011), and whole brain data were segmented in five tissue classes (cortical grey matter, sub-cortical grey matter, white matter, cerebrospinal fluid (CSF) and pathological tissue) using FSL FAST (Zhang et al., 2001) which was used for the purposes of an anatomically constrained tractography. Cortical surface segmentation of white matter and pial surfaces were generated from these processed scans using Freesurfer 6.0 (Fischl, 2012) and were manually assessed and quality controlled.

Resting-state fMRI preprocessing involved dropping the first 5 TRs, image reorientation to LPI and motion correction (registering all timepoints volumes to the mean image, rigid body transformation 3dvolreg). Motion outliers (i.e., motion spikes) were estimated using *fsl_motion_*outliers. We then performed distortion correction, applied a high-pass filter removing frequencies below 0.01Hz. Furthermore, we also applied additional preprocessing outside of micapipe on the rs-fMRI volume to ensure optimal compatibility with SPARK. This included the regression of motion spike outliers and CSF and WM nuisance variables estimated during the micapipe processing. We also applied temporal averaging of every three timepoints to create a high signal-to-noise (SNR) while reducing the computational load of the BOLD volumes, allowing us to apply SPARK within a similar dimensionality as in our previous investigations (Lee et al., 2016, 2018). A transformation matrix first estimating the co-registration of the BOLD volumes to the subject native T1w anatomical space and then to a 4mm isotropic voxel version of the ICBM 2009c Nonlinear symmetric template using FSL’s *flirt* (Jenkinson and Smith, 2001; Jenkinson et al., 2002). Co-registration was estimated by maximizing mutual information between volumes as the cost function. Finally, spatial smoothing with a full-width-half-maximum (FWHM) of 8mm was applied constrained within the boundaries of a binary grey matter mask using *mri_fwhm* from Freesurfer.

Diffusion-weighted imaging was preprocessed using MRtrix3 (Tournier et al., 2019). Preprocessing included denoising, Gibbs ringing removal, b0 intensity normalization, brain masking, and correction for susceptibility distortion, eddy currents, and head motion. Gradient tables were verified and corrected, and a bias field correction was applied. All DWI scans were aligned using a rigid-body co-registration. ADC and FA maps were derived from the diffusion tensor image. ADC is a visual representation of the molecular diffusion rate of a voxel, where lower values represent lower mean diffusivity, and FA is a measure of the degree of anisotropic diffusion found within the voxel ranging from 0 (isotropic diffusion) to 1 (anisotropic diffusion) (Soares et al., 2013).

### Connector Hub Estimation using Sparsity-based analysis of reliable *k*-hubness (SPARK)

SParsity-based Analysis of Reliable *k-*hubness (SPARK), is a sparse coding method designed to specifically assess the discrete spatial overlap between resting state networks at a voxel level. This method has been extensively outlined in previous work from our group. (Lee et al., 2016, 2018, 2022, 2026). Briefly, SPARK aims to explain the rs-fMRI BOLD time series for each voxel as a linear combination of multiple RSN time series using: (1) a sparse single value decomposition (SVD) of the data to estimate a sparse code for each voxel and (2) a circular bootstrap resampling strategy to assess reliability of SPARK results. The core of the SPARK is a *k-*SVD algorithm (Aharon et al., 2006), which is a sparse dictionary learning algorithm which first estimates *N*-atom time courses from the data before assessing the sparse decomposition of these time courses in each voxel. This results in a maximum of *k* interacting time courses characterizing the BOLD signal within one voxel. The number of non-zero entries of this sparse General Linear Model (Lee et al., 2011), also known as sparse code, denotes our hubness measure that we will refer to as *k-*hubness. Each atom time course is associated to an atom spatial map, denoting a consistent RSN. Every RSN estimated using k-SVD is therefore allowing “sparse” spatial overlap between RSNs by construction. The total number of atoms (*N*) was automatically estimated for each subject using an minimum mean description (MDL)-based approach, by testing *N* values from 10–40 and selecting the value yielding the MDL across sparsity levels (Lee et al., 2018). In a second step, to assess a reliable decomposition of *k-*hubness, *k-*SVD was applied sequentially on 200 bootstrap samples extracted from the original data using a circular bootstrapping approach, as proposed in Bellec et al., 2010. To select the most reliable k-SVD and their *k-*hubness estimations, a hierarchical clustering was applied over all atom maps across all bootstraps, resulting in *N* reliable atom maps (averages within the *N* resulting clusters) and the corresponding sparse coding matrix. Less reproducible elements, characterized by low signal amplitudes relative to the background noise, were then considered as background noise assuming a Gaussian distribution and were subsequently removed from the sparse coding matrix using a threshold of *p* < 0.05. Resulting reliable atom maps were then manually labeled as a RSN or noise (e.g.: cardiac, respiration). The identified noisy atom maps were removed for the subsequent analysis, and the remaining atom maps representing the canonical RSNs were used to generate a final *k-*hubness map by counting the number of non-zero coefficients in each column of the sparse code matrix, i.e.: the number of overlapping RSNs within each voxel. *k-*values typically range between 1 to 10 networks per voxel. Within the SPARK framework, *hubness* assesses the propensity of a voxel to be associated with an RSN. Each voxel is assigned a discrete *k-*hubness value, where *k* represents the number of RSNs from whom time courses are interacting within this voxel, denoting spatial overlap between networks. A *k-*hubness value of 1 indicates that the voxel is associated with one RSN, whereas a *k-*hubness value larger than one implies the voxel is associated with many networks and is a hub. SPARK offers the unique possibility to identify exactly what are the underlying RSNs associated with each hub. Finally, a *k-*hubness value of 0 is indicative of an unreliable finding over bootstrap samples. We applied the SPARK method to the preprocessed rs-fMRI BOLD time series of each subject, allowing us to estimate the organization of functional hubs across individual brain networks at a voxel level.

### Data visualization

Using tools from Connectome Workbench (https://github.com/Washington-University/workbench), finalized *k-*hubness maps estimated in ICBM 2009c Nonlinear symmetric template (4mm resolution), and ADC and FA volumetric maps estimated in DWI-space were mapped to the native surface using *volume-to-surface-mapping* using the *enclosing voxel* mapping method. These maps were then resampled to the fsLR-5k surface template provided by micapipe using *metric-resample*. Finally, a 10mm smoothing was applied along the surface using *metric-smoothing*. Visualization of cortical maps was performed using *plotSurfaceROIBoundary* (Oldham, 2026). The hippocampal surfaces were generated using HippUnfold v1.3.0 (DeKraker et al., 2022), allowing for vertex-wise comparisons between various maps visualized using hippomaps v0.1.0 (DeKraker et al., 2025), allowing us to study the hippocampus at a finer spatial scale given the pathological relevance in TLE. Hippocampal maps were projected to a 2mm density template surface using *volume-to-surface-mapping* and underwent a 3mm smoothing kernel along the surface using *metric-smoothing* from workbench. Finally, for our subcortical structure analysis, we computed average values for each element of the subcortical segmentation provided by Freesurfer (bilateral accumbens, amygdala, caudate, hippocampus, pallidum, putamen, and thalamus) and visualized these regions using ENIGMA toolbox (Larivière et al., 2021)

### Patients versus controls *k-*hubness alterations in temporal lobe epilepsy

Vertex-wise *k-*hubness maps in patients were first Z-scored relative to healthy controls and sorted ipsilateral/contralateral to the epilepsy focus, allowing us to increase the power of our analysis by lateralizing the epilepsy foci. We fit a surface-based fixed effect linear model to study changes in *k-*hubness values between patients and controls using Brainstat v0.2.4 (Larivière et al., 2023). Controlling for age and sex, we defined the following linear regression model for each vertex *i*:

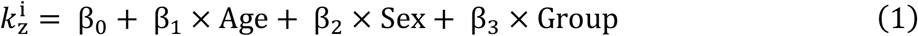

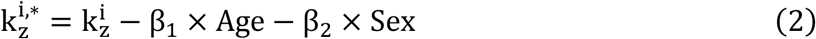

From this model, we obtain 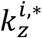 which are the 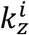Z-scored *k-*hubness maps corrected for age-and-sex for each subject. The region exhibiting significant differences between 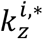 values between our patient cohort and controls was estimated using a two-sided t-test. Findings were corrected for multiple comparisons using Random Field Theory along the cortical surface and unfolded hippocampus surface and were corrected using False Discovery Rate (FDR) in subcortical structures. We define 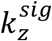 as the resulting significant clusters obtained from the contrast between the epilepsy and healthy control groups (*β*_3_) while correcting for the fixed effects of age (*β*_1_) and sex (*β*_2_).

### Relationship between SPARK networks and significant clusters

During SPARK, each subject underwent the manual identification and labeling of every atom map estimated by the algorithm. Using the labeled maps across healthy controls, we generated a SPARK data-driven average resting state network maps for the auditory, cerebellum, default mode, dorsal attention, executive control, hippocampus, precuneus, salience, sensorimotor, basal ganglia, temporal lobe, visual lateral, visual medial visual occipital, and visuospatial networks, resulting in a dataset-specific resting state network atlas (SPARK_RSN_). The naming of the RSNs were inspired by the typical conventions found across the literature (Heine et al., 2012; Di and Biswal, 2014; Lv et al., 2018). To generate SPARK_RSN_, the subject-level atom maps were initially mapped along the fsLR-5k cortical surface and underwent binarization (1 = vertex was actively a part of the atom map, 0 = vertex was not a part of the atom map). Then for each network, the corresponding binarized atom maps identified over the whole population of controls were averaged at every vertex with a threshold of 20% being applied to each vertex. The threshold allowed us to only keep the vertices that where at a minimum present in 20% of the controls, while minimizing the effect of noisy findings. Subcortical contribution was computed in the same manner, but in volumetric space. Finally, we then used the SPARK_RSN_ atlas to compute the involvement of the significant cluster exhibiting differences between patients and controls, 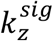 to each network of SPARK_RSN_ in the following manner: area of vertices involved in overlap

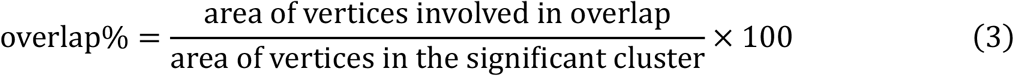

### Hub Disruption Index

The hub disruption index (HDI) proposed by (Achard et al., 2012; Lee et al., 2018), is a group level metric quantifying how hubs in a region of interest change in pathological conditions when compared to the distribution of hubness in a healthy cohort. We adapted the originally proposed metric to fit within the framework of our methodology. To do so, we first compute the variable 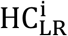, the normalized mean of the hemispheric average at each vertex for every healthy subject to reduce the bias of the ipsilateral/contralateral flipping performed on our epilepsy cohort.

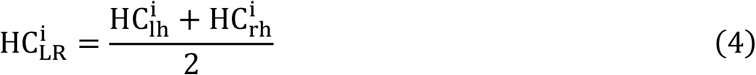

where 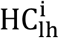 and 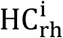 are the 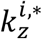 Z-scored values measured within the left and right hemispheres respectively for each healthy control. We can then compute our updated hub disruption metric as:

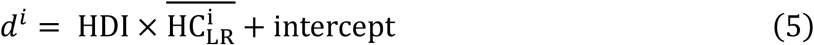

Where the y-axis (*d*^*i*^) is the Cohen’s *d* value at each vertex *i*:

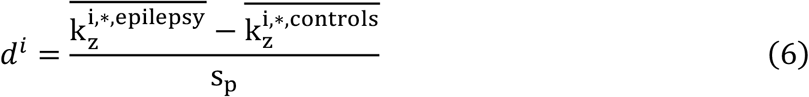

where we define 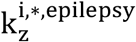 as the age-and-sex corrected Z-score maps of the TLE patients, and 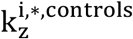 are the age-and-sex corrected Z-score maps of the healthy control cohort. We further define *s*_*p*_ as the pooled standard deviation:

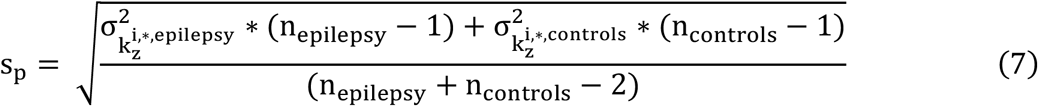

where 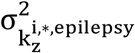 is the standard deviation of the age-and-sex corrected Z-score maps across TLE patients, and 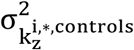 is the standard deviation of the age-and-sex corrected Z-score maps across healthy subjects. Finally, we define 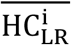 is the average value of 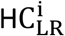 across all healthy controls, which we denoted as the normalized mean of hemispheric average of controls. Therefore, HDI and intercept can be estimated by applying the linear model of equation (5) for all vertices of a specific region of interest.

In our updated model, the hub disruption index, reflected as the slope of the linear regression model represents the rate of degradation (in the case of a negative slope) or strengthening (in the case of a positive slope) of *k-*hubness relative to the hemispheric average 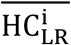. A degradation represents the disruption of connections whereas a strengthening represents an emergence of new connections. The steeper the slope, the quicker pathological hubs are changing relative to hubs in healthy controls. The y-intercept represents the Cohen’s *d* value of 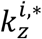 at the mean hemispheric averaged Z-scored values over the cluster, where a negative intercept represents an initial value of hub disruption, and a positive intercept represents and initial value of hub emergence. We perform this analysis in the 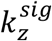 clusters which were found along the cortical surface, along the unfolded hippocampus surface, and within subcortical structures (volumetric analysis) and their contralateral homologous regions.

To assess the significance of the values of the slope (HDI) and intercept, we perform three types of significance testing depending on the region. Significance along the cortical surface was performed using spin testing incorporated within the BrainSpace toolbox v0.1.4 (Vos de Wael et al., 2020), where the y-axis data (i.e.: Cohen’s *d* map) was permutated N_permutations_ = 1000 times to generate surrogate maps preserving the spatial autocorrelation of the data. The slope and intercept were then computed between each surrogate map and 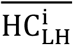 to generate a null disruption for each value respectively. Depending on whether our actual computed value of HDI or the intercept (i.e.: trueValue) was positive or negative, we computed a one-sided non-parametric test to assess the corresponding *p*-value using the null distribution (null) and the number of permutations (N_permutations_) generated using spin test:

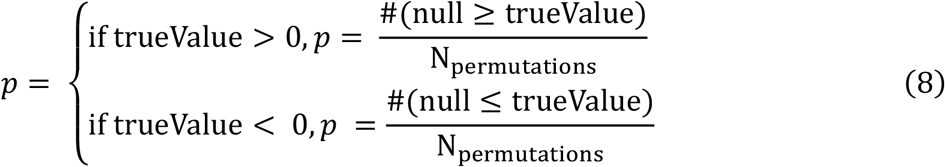

Significance testing on the hippocampal ROIs was performed using spin testing using the *hippomaps*.*stats*.*spin_test* function integrated within the hippomaps software (DeKraker et al., 2025) to generate the surrogate maps. The same methodology follows for computing a corresponding *p-*value for the slope and intercept. For all methods, *p < 0*.*05* was considered significant.

Significance testing on subcortical ROIs was performed in volumetric space by using Moran’s Randomization which is integrated within the BrainSpace toolbox v0.1.4 (Vos de Wael et al., 2020) to generate the surrogate maps. Here, we masked the region of interest and built a 6-neighborhood adjacency matrix with

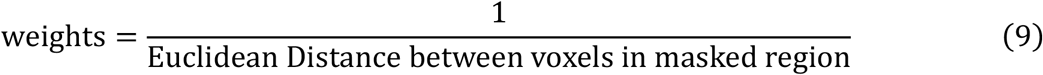

and computed the Moran eigenvectors to generate the Cohen’s *d* surrogate maps in the ICBM 4mm space. From here, the same methodology follows for computing a corresponding *p-*value for the slope and intercept.

### Relationship between hubness and whole-brain microstructure

To study changes in white matter microstructure, vertex-wise ADC and FA maps were Z-scored relative to controls and were sorted ipsilateral/contralateral to the epilepsy focus before applying an identical surface based-linear model to one used for *k-*hubness, creating age-and-sex corrected Z-score maps for FA 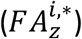 and ADC 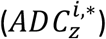, along with their respective significant clusters 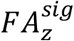 and 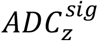 exhibiting differences between patients and controls. To study the relationship between the microstructure and *k-*hubness we performed three analyses: (i) we recomputed 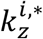 in a hierarchical manner, first regressing out the effects of ADC and FA at each vertex across the entire population, followed by regressing out age and sex from the corrected values, as in equation (1), resulting in 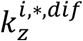. We then recomputed the Cohen’s *d* and performed a *t*-test to re-evaluate whether there were changes in the structure of 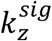; (ii) performed a patient-specific *k-*hubness profile analysis, where we compared the average 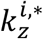 and 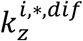 values within 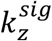 for each subject, and (iii) computed the correlation coefficient (*r*) between 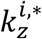 and 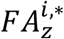 and 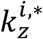 and 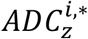 in four regions of interest: 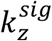 and the homologous contralateral region along the cortical surface, and within the entire ipsilateral and contralateral unfolded hippocampal surface. This analysis was not performed for any subcortical structures. Significance was assessed using spin permutation tests, with a significance threshold of *p* < 0.05 (uncorrected).

### Relationship between hubness, white matter microstructure and seizure freedom

We finally evaluated the trifold relationship between pre-operative *k-*hubness and white matter microstructure with post-operative seizure freedom. This analysis was applied on a subgroup of 9 seizure-free patients who underwent surgical intervention after their initial imaging, and for whom postoperative follow-up after one year has been obtained and classified as seizure free (Engel IA) (Epilepsy (ILAE) 1997–2001 et al., 2001). At a group level, we first computed 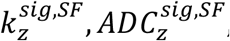, *and* 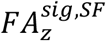 for the 9 seizure free patients. Then, at a subject level, we assessed both (i) the *k-*hubness and diffusion metric linear relationship and (ii) the *k-*hubness and hippocampal volume linear relationship, before also including non-seizure free (n=3) and unoperated patients on the resulting linear fits. Hippocampal volume was computed as the sum of the volumes of the hippocampal subfields estimated by HipUnfold. Volumes were then adjusted for head size by regressing out the effect of intracranial volume, as estimated by FreeSurfer, prior to subsequent analyses (Voevodskaya et al., 2014).

## Acknowledgements

**A.D**. was funded by Quebec Bio-Imaging Network and McGill Medicine. **E.D**. was funded by the Quebec Sleep Research Network. **A.N**. was funded by CIHR and FRQ-S. **J.C**. was funded by the Vanier Canada Graduate Scholarship. **E.S**. acknowledges funding from the Vanier Canada Graduate Scholarship. **T.A** is funded by Savoy Foundation Postdoctoral Fellowship. **C.A**. was supported by the doctoral award of Savoy Foundation, Fonds de Recherche du Québec – Santé, and Canadian Institutes of Health Research (CIHR). **J.R**. is funded by a Banting Postdoctoral Fellowship. **R.R.C**. is funded by the Fonds de Recherche du Québec—Santé (FRQ-S). **K.L**. was supported by the Wallace H. Coulter Foundation. **B.C.B**. acknowledges support from the Canadian Institutes of Health Research, CIHR (FDN-154298, PJT-174995, PJT-206196, CIHR PJT-203761, CIHR PJT-191853), SickKids Foundation (NI17-039), Natural Sciences and Engineering Research Council (NSERC RGPIN-2025-05932), Azrieli Center for Autism Research of the Montreal Neurological Institute (ACAR), BrainCanada, FRQ-S, the Helmholtz International BigBrain Analytics and Learning Laboratory (Hiball), Healthy Brains and Healthy Lives (HBHL), Centre for Aging + Brain Health Innovation (CABHI), the Canada Research Chairs Program (CRC), and the Centre of Excellence in Epilepsy at the Neuro (CEEN). **C.G**. acknowledges support by Canadian Institutes of Health Research, CIHR (PJT-159448, PJT-195828), the Natural Sciences and Engineering Research Council of Canada (NSERC) Discovery grant (RGPIN 2025 06816), and a research team grant from Fonds de Recherche du Québec—Nature et Technologie (FRQ-NT).

## CRediT authorship contribution statement

**Arielle Dascal**: Conceptualization, Methodology, Software, Validation, Investigation, Data curation, Writing – Original Draft, Visualization. **Édouard Delaire**: Software, Writing - Review & Editing, Visualization. **Alexander Ngo:** Software, Data curation, Writing - Review & Editing. **Judy Chen:** Software, Data curation, Writing-Review & Editing. **Ella Sahlas:** Data Curation, Writing – Review & Editing. **Thaera Arafat:** Data curation. **Chifaou Abdallah:** Writing – Review & Editing. **Mahdi Mobarak-Abadi**: Writing – Review & Editing. **Jessica Royer:** Data Curation, Writing-Review & Editing. **Raul Rodriguez Cruces:** Data Curation, Writing – Review & Editing. **Neda Bernasconi:** Data Curation. **Andrea Bernasconi:** Data Curation. **Raluca Pana:** Data Curation. **Kangjoo Lee**: Software, Methodology, Writing – Review & Editing. **Boris C. Bernhardt:** Conceptualization, Methodology, Data Curation, Software, Writing – Review & Editing, Supervision, Project Administration, Funding Acquisition. **Christophe Grova:** Conceptualization, Methodology, Software, Writing – Review & Editing, Supervision, Project Administration, Funding Acquisition.

